# The transcriptome-wide landscape and modalities of EJC binding in adult *Drosophila*

**DOI:** 10.1101/459354

**Authors:** Ales Obrdlik, Gen Lin, Nejc Haberman, Jernej Ule, Anne Ephrussi

## Abstract

Splicing-dependent assembly of the exon junction complex (EJC) at canonical sites −20 to −24 nucleotides upstream of exon-exon junctions in mRNAs occurs in all higher eukaryotes and affects most major regulatory events in the life of a transcript. In mammalian cell cytoplasm, EJC is essential for efficient RNA surveillance, while in *Drosophila* the most essential cytoplasmic EJC function is in localization of *oskar* mRNA. Here we developed a method for <u>is</u>olation of <u>p</u>rotein complexes and <u>a</u>ssociated <u>R</u>NA-<u>t</u>argets (ipaRt), which provides a transcriptome-wide view of RNA binding sites of the fully assembled EJC in adult *Drosophila*. We find that EJC binds at canonical positions, with highest occupancy on mRNAs from genes comprising multiple splice sites and long introns. Moreover, the occupancy is highest at junctions adjacent to strong splice sites, CG-rich hexamers and RNA structures. These modalities have not been identified by previous studies in mammals, where more binding was seen at non-canonical positions. The most highly occupied transcripts in *Drosophila* have increased tendency to be maternally localized, and are more likely to derive from genes involved in differentiation or development. Taken together, we identify the RNA modalities that specify EJC assembly in *Drosophila* on a biologically coherent set of transcripts.

## Introduction

The exon junction complex (EJC) consists of a hetero-tetramer core composed of eIF4AIII, Mago, Y14 and Barentsz (Btz) (1,2), and auxiliary factors that form the EJC periphery (3). The complex assembles on mRNAs during splicing, −20 to −24 nucleotides (nt) upstream of exon-exon junctions (4). EJC assembly is a multi-step process that begins with the CWC22-mediated deposition of the DEAD-box helicase eIF4AIII on nascent pre-mRNAs (5-7) and is followed by the recruitment of Mago and Y14, forming a pre-EJC intermediate. The pre-EJC is stably bound to RNA due to the ATPase inhibiting activity of the (non-RNA-binding) Mago-Y14 heterodimer, which “locks” eIF4AIII helicase in its RNA-bound state (1,2,8-10). Once formed, the pre-EJC is completed by recruitment of Barentsz (Btz), forming mature EJCs (1,3,11). The roles of the EJC in post-transcriptional control of gene expression are manifold. In the nucleus, EJC subunits have been shown to play a role in splicing (12-15), mRNA export (16), and nuclear retention of intron-containing RNAs (17). In the cytoplasm, the EJC is reported to play a role in translation (18,19), nonsense-mediated decay (NMD) (7,20-26) and RNA localization (23,27-30). While most of the EJC’s functions appear conserved, in *Drosophila* the EJC is not crucial for NMD (31), but it is essential for *oskar* mRNA transport and localization within the developing oocyte (23,27-30,32,33). To better understand the engagement of the EJC in the fly we set out to define the EJC mRNA interactome in adult *Drosophila melanogaster* using a novel strategy that stabilizes mRNA binding proteins (mRBPs) associated with their RNA templates within multi-protein mRNP assemblies. This method through the use of the crosslinking agent dithio(bis-) succinimidylpropionate (DSP) captures stable and transient protein interactions in close proximity (34,35) and allows definition of the binding sites of specific protein (holo-) complexes associated with their RNA templates (ipaRt). Indeed, our analysis of EJC protected sites defined by ipaRt reveals that, in *Drosophila*, EJC binding occurs at canonical deposition sites (4), with a median coordinate ∽22 nt upstream of exon-exon junctions. While in mammals EJC mediated protection outside canonical sides has been reported (36,37), we find that in *Drosophila* the degree of non-canonical EJC mediated RNA protection is minimal. We show in *Drosophila* that RNA polymerase II transcripts primarily protected by the EJC derive from genes involved in differentiation or development, while mRNAs primarily protected by mRBPs derive from genes with homeostatic functions. Our analysis suggests that the EJC’s bias for transcripts is in *Drosophila* a consequence of several modalities in the genes’ architecture, particularly the number of splice sites and intron length. Moreover, EJC binding is enhanced by adjacent RNA secondary structures and CUG-rich hexamer sequences located 3’ to the EJC binding site. These modalities have not been identified in previous studies of mammalian EJC binding sites (36-38), which could either reflect the greater specificity of our method for the fully assembled EJC, or could reflect differences in EJC binding between flies and human. Our study provides a first comprehensive transcriptome-wide view of EJC-RNA interactions in a whole organism, which unravels RNA modalities that contribute to the unforeseen biological coherence of the bound transcripts.

## Results

### Stabilization of the Exon Junction Complex on mRNAs by DSP

The EJC is maintained in its RNA-bound state through the direct interaction of the Mago-Y14 hetero-dimer with the otherwise dynamically binding RNA helicase eIF4AIII (1-3,9,22,39). EJC binding to RNA is labile, as under stringent washing conditions (∽ 1M salt concentrations) the interaction of eIF4AIII and Mago-Y14 is abolished and the RNA is released from the complex (36). We therefore hypothesized that the EJC might be stabilized on its RNA targets by introducing covalent bonds between the Mago-Y14 heterodimer and eIF4AIII, which would render the protein-RNA complex resistant to the high salt concentrations commonly used in iCLIP studies. Furthermore, a stabilized EJC complex would enable us to “pull” on EJC subunits other than the RNA-binding eIF4AIII, ensuring isolation of the complex under stringent conditions. To test this we made use of the bivalent crosslinking agent dithio(bis-) succinimidylpropionate (DSP), which crosslinks reversibly primary amino groups of polypeptides in close proximity (34,35). We isolated poly(A) containing mRNPs on an oligo d(T)_25_ resin (40,41) from cytoplasm of adult *Drosophila* either kept untreated or treated with UV, DSP, or UV plus DSP (Figure 1). SDS-PAGE silver staining and western analyses of mRNA-RNP precipitates (Figure 1A,B) revealed that irradiation of cytoplasmic lysates by UV *ex vivo* only marginally increased co-precipitation of proteins with poly(A) containing RNAs (Figure 1A, lane 7 and 8). Upon UV irradiation, only faint signals of known RBPs such as eIF4AIII and cytoplasmic poly(A) binding protein (PABP) were detected in the poly(A) RNA precipitates. Non RNA-binding EJC subunits such as Y14 were not detected (Figure 1B, lane 7 and 8, Figure 1C) in agreement with previous observations (41). In contrast, treatment of cytoplasmic lysates with DSP led to strong protein co-precipitation with mRNAs (Figure 1A, compare lanes 7-10). Western analysis of precipitates from DSP and UV-DSP treated cytoplasmic lysates revealed strong signals not only for direct mRNA binding proteins such as eIF4AIII and PABP, but also mRNP components not directly bound to RNA, such as Y14 (compare Figure 1A-B, lanes 7-10, Figure 1C). In none of the precipitates were cytoplasmic proteins such as kinesin heavy chain (Khc) or the small ribosomal subunit protein RpS6 observed (Figure 1B, lanes 7-10, Figure 1C), confirming the stringency of the assay. Furthermore, precipitates from the “beads” only control were free of all proteins tested (Figure 1A, lane 6), indicating that artifacts due to DSP or UV derived treatment are unlikely. These observations suggest that, for EJC stabilization on RNA, DSP-mediated covalent bond formation between individual EJC subunits is superior to UV-crosslinking and support the use of this agent when studying other mRNP assemblies (Figure 1D).

**Figure 1.**
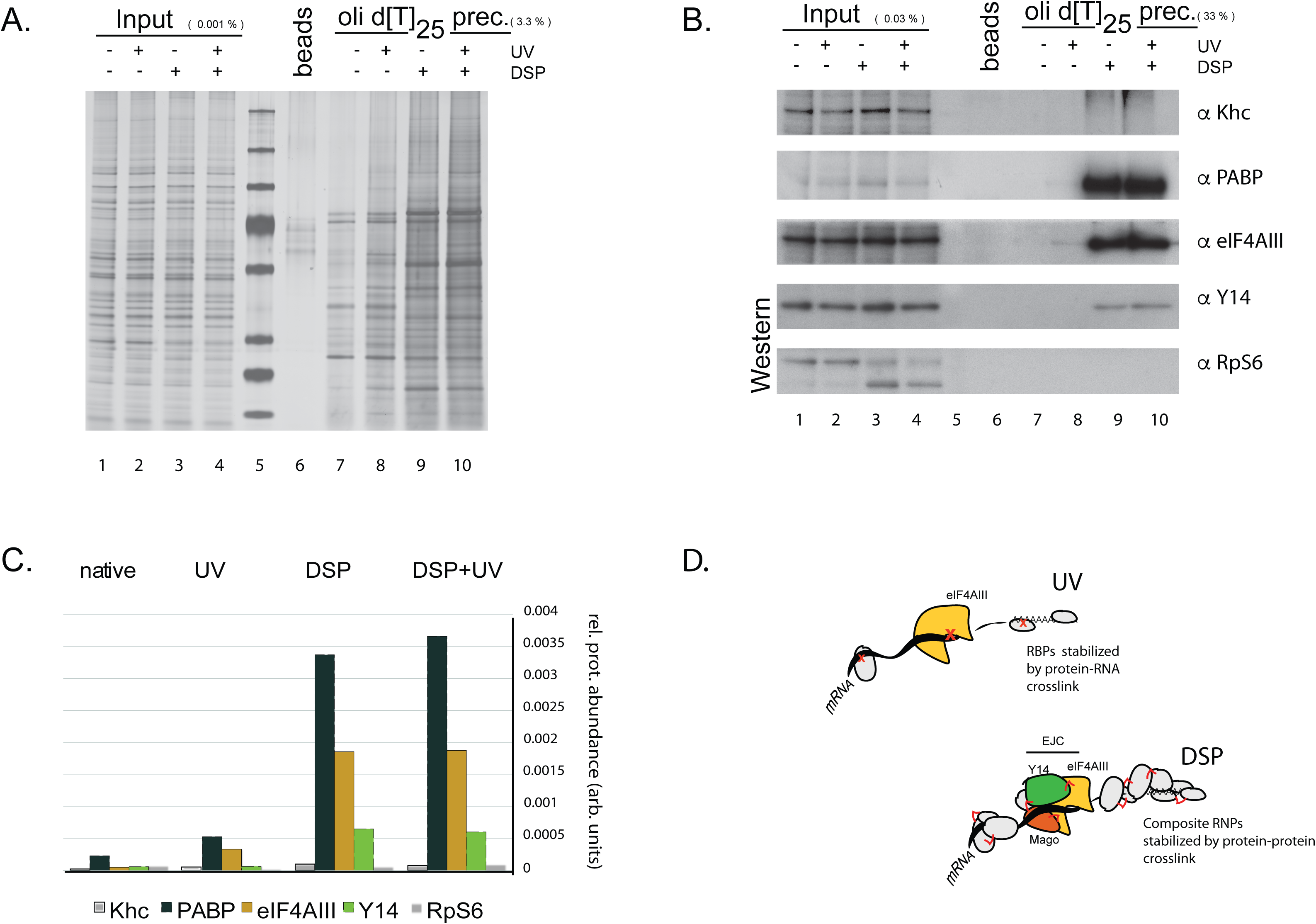
**Dithio(bi-)succinimidyl-propionate (DSP), stabilizes EJC in its mRNA bound state.** **(A) SDS-PAGE and silver staining of proteins co-precipitated with oligod(T)_25_ bound mRNAs from cytoplasm.** Left to right: Input cytoplasmic samples (0.01% of total input; lanes 1-4); protein MW standards (lane 5); “beads-only” control precipitate from “all-condition-mixture” (3.3%, lane 6); individual oligo-d(T)_25_ precipitates (3.3%; lanes 7-10). 931 Treatment conditions,“untreated”, “UV irradiated”, “DSP supplemented” and “UV irradiated-DSP supplemented” are indicated (+ or -) above image. Note that for samples treated with DSP *ex vivo*, UV-exposure does not increase the amount of recovered proteins. **(B) Western blot analysis of oligo-d(T)_25_ precipitates.** Gel loading order same as in (A); 0.03% of the input cytoplasm and 33% of each precipitate were resolved. Blot was probed with antibodies directed against the proteins indicated at the right of the panel. Khc: kinesin heavy chain; PABP: cytoplasmic poly A binding protein; RpS6 = small ribosomal subunit protein S6. **(C) Quantification of proteins detected on the western blot.** Crosslinking conditions are indicated in upper panel. Signals were quantified by densitometry measurement using the Fiji image analysis package. Relative protein abundance is the fraction of the signal in the precipitate relative to the cytoplasm. Note that in contrast to the RNA-binding eIF4AIII, the non-RNAbinding EJC subunit Y14 is only detected when the cytoplasm was treated with DSP. **(D) Schematic of the net effect of crosslinking with UV versus DSP.** Exposure of the cytoplasm to UV leads primarily to stabilization of direct protein-RNA interactions. DSP treatment results in efficient retention of proteins associated with RNA by stabilization of polypeptide interactions either within individual an RBP or between an RBP and other moieties within a complex.

### ipaRt: a novel approach for high quality isolation of EJC complexes associated to RNA templates

Of all the tagged EJC subunits we tested in the fly, GFP-Mago showed the highest degree of incorporation into endogenous EJCs (Figure S1B). Furthermore, the eIF4AIII subunit was additionally found to co-sediment with polysome fractions in sucrose density gradients, independently of Mago-Y14 (Figure S1C), suggesting that the DEAD-box helicase might have yet unknown EJC independent functions in the fly (see Supplementary Results). Therefore, we carried out EJC specific RNA immunoprecipitation (RIP) from DSP treated cytoplasmic extracts prepared from GFP-Mago and GFP-tag expressing flies. By titrating salt and detergent concentrations, we identified stringent washing conditions (see Materials and Methods) that yielded high quality RNA profiles from GFP-Mago RIPs and only scant RNA profiles from GFP control RIPs, compared with standard IP washing conditions (Figure 2A; compare lanes 2-5). To test whether the presence of RNA in GFP-Mago precipitates was a consequence of its incorporation into the EJC rather than by virtue of transient interactions of Mago with other RBPs, we treated the DSP treated or untreated lysates immunoprecipitates with RNAseI. Western analysis revealed the presence of all tested EJC subunits in the GFP-Mago precipitates (Figure 2B, lanes 7-8), but no protein other than GFP itself in the GFP only controls (Figure 2B, lanes 3-4). In contrast we detected no signal for the proteins probed in the “beads only” control precipitates (Figure 2B, lanes 2 and 6) and observed signals for the RNA non-binding Khc only in lysate inputs (Figure 2B, compare lanes 1, 5; lanes 2-4, 6-8), indicating high stringency of the assay.

**Figure 2.**
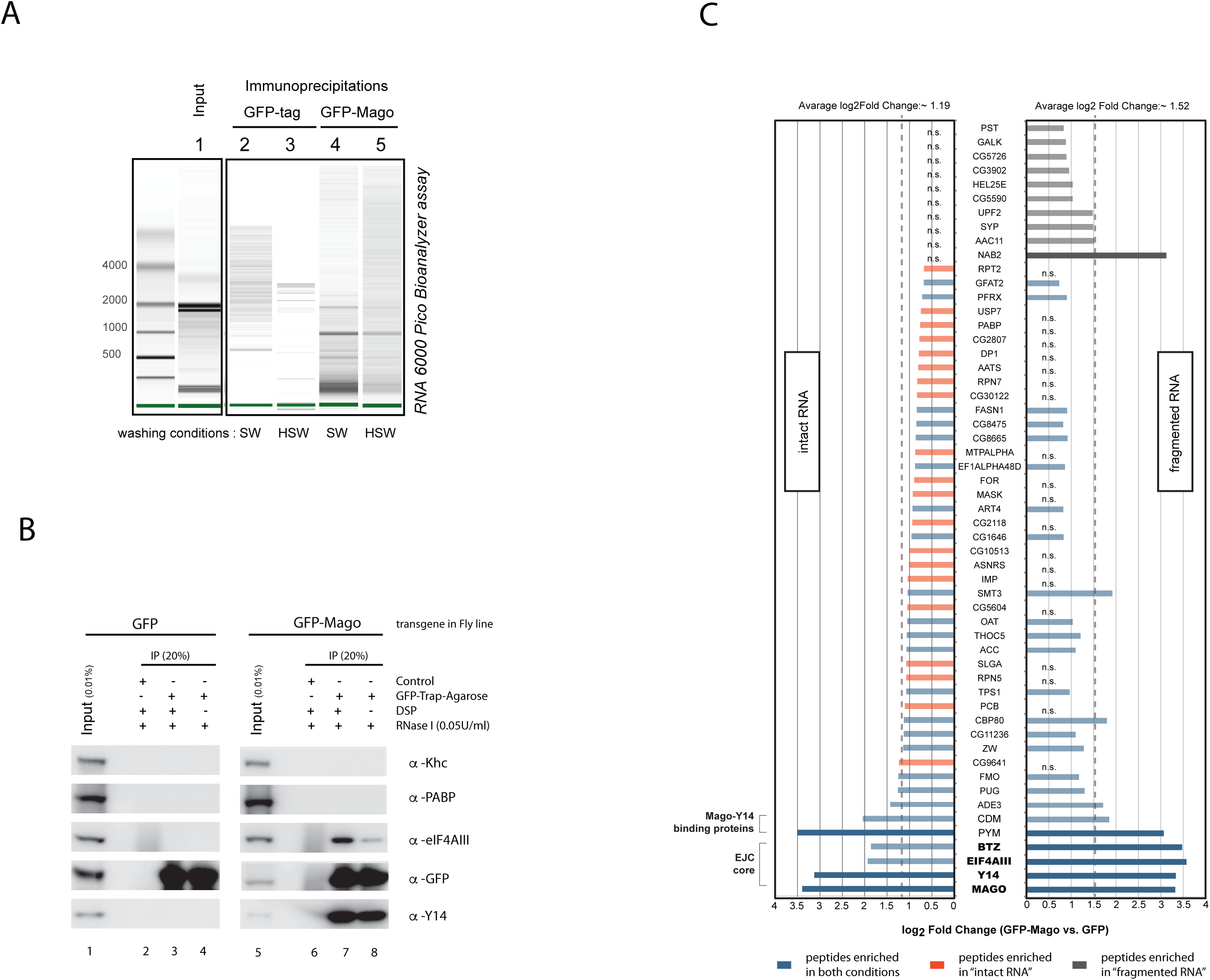
DSP crosslinking stabilizes EJC for stringent immunoprecipitation and RNA fragmentation. **(A) Comparison of IP washing conditions for RNA isolation.** Anti-GFP RNA IP assays on DSP-treated cytoplasm of GFP-Mago and GFP tag expressing flies. 0.02% of RNA isolated from input lysate (lane 1) and 20% from GFP tag (lanes 2 and 3) and GFP-Mago (lanes 4 and 5) RIP precipitates were resolved by capillary gel electrophoresis on an RNA 6000 Pico Chip Bioanalyzer. Washing conditions are indicated at bottom. SW: standard stringency washing conditions; HSW: high stringency washing conditions (see Materials and Methods). **(B) DSP stabilized EJC core is resistant to RNaseI treatment.** Effects of RNaseI treatment on GFP-Mago co-immunoprecipitations in DSP crosslinked and untreated cytoplasm. Western blots of anti-GFP IPs from DSP treated and from untreated cytoplasm of GFP-Mago (lanes 5-8) and GFP tag (lanes 1-4) expressing flies processed under HSW conditions. Primary antibodies utilized are indicated on the right. Inputs (0.01%) are shown in lane 1 and 5. Bead-only control precipitates are shown in lanes 2 and 6. IP precipitates (20%) from DSP treated (lanes 3 and 7) or untreated cytoplasm (lanes 4 and 8). Details of samples and experimental conditions indicated at top of panel. **(C) RNA fragmentation depletes EJC unrelated proteins from GFP-Mago co-precipitates.** Anti-GFP IPs (HSW condition) from DSP treated cytoplasm of GFP-Mago and GFP tag expressing flies. Precipitates were subjected to isobar labeling and peptide content was defined by tandem mass spectrometry. Presented bar plots of protein-enrichment are defined by Limma(80). GFP Mago specific protein enrichments in “intact RNA ” and “fragmented RNA ” conditions are highlighted in left and right plot, respectively. Y-axis shows individual proteins detected. X-axis shows scale of enrichment (Log2 fold change). Dashed line indicates average enrichment of all significant proteins in each condition. Protein-enrichment >2x or <2x average-enrichment is indicated by solid or transparent bars, respectively. Enrichment bar color legend is highlighted at the bottom. ns: non-significant (p.adj.>0.05).

The stabilizing effect of DSP on the EJC is evident from the enhanced eIF4AIII signals in GFP-Mago precipitates when cytoplasm was treated with DSP prior to immunoprecipitation (Figure 2B, lanes 5, 7-8). Conversely, the PABP signal in the GFP-Mago precipitates disappeared upon incubation with RNase of both the DSP treated and the untreated samples (Figure 2B, lanes 1, 5; lanes 2-4, 6-8). This shows that even when exposed to DSP, proteins whose association with the EJC is bridged by RNA can be removed by RNA fragmentation.

To confirm the “cleansing effect” of RNAseI, we perfomed IPs from DSP treated cytoplasm with or without an RNA fragmentation step and analyzed the precipitates by tandem mass spectrometry (MS). Expression set analysis of MS signals obtained in GFP-Mago and GFP control precipitates identified 45 versus 35 significantly enriched proteins in the RNase untreated and treated samples respectively (Figure 2C, Figure S3). While all EJC subunits were enriched independently of RNA integrity (Figure 2C), only upon RNA fragmentation was Btz enriched to a similar degree as Mago, Y14, and eIF4AIII. Except for the poly(A) binding protein Nab2 (42), no EJC-unrelated RBPs showed significant enrichment upon RNA fragmentation (Figure 2C), showing that RNA fragmentation by RNaseI increases the sensitivity and specificity of EJC IPs.

Based on these findings, we conclude that DSP is a suitable tool for stablilization and isolation of EJC RNA complexes from animal tissues and that our protocol provides a reliable approach for isolation of proteins (or protein complexes) associated to their RNA targets. Consequently we termed our experimental strategy “ipaRt”.

### EJC binding in *Drosophila* cytoplasm maps to canonical deposition sites

We sought to determine which sites in the *Drosophila* transcriptome are protected by the EJC as opposed to other mRNA binding proteins (mRBPs). To this end, we performed EJC ipaRt and oligo(dT) mediated mRNP capture in parallel, both followed by an RNase digestion step (see Material and Methods), and constructed cDNA libraries of the protein-protected RNA fragments as previously described for iCLIP (43). Analysis of the sequencing results revealed that more than 92% of all reads aligned uniquely to the *Drosophila* genome (Figure S3B). 85% of EJC ipaRt reads mapped to exons as opposed to 34% in the mRBP footprinting mapped reads, indicating specificity of the ipaRt library (Figure 3A).

**Figure 3.**
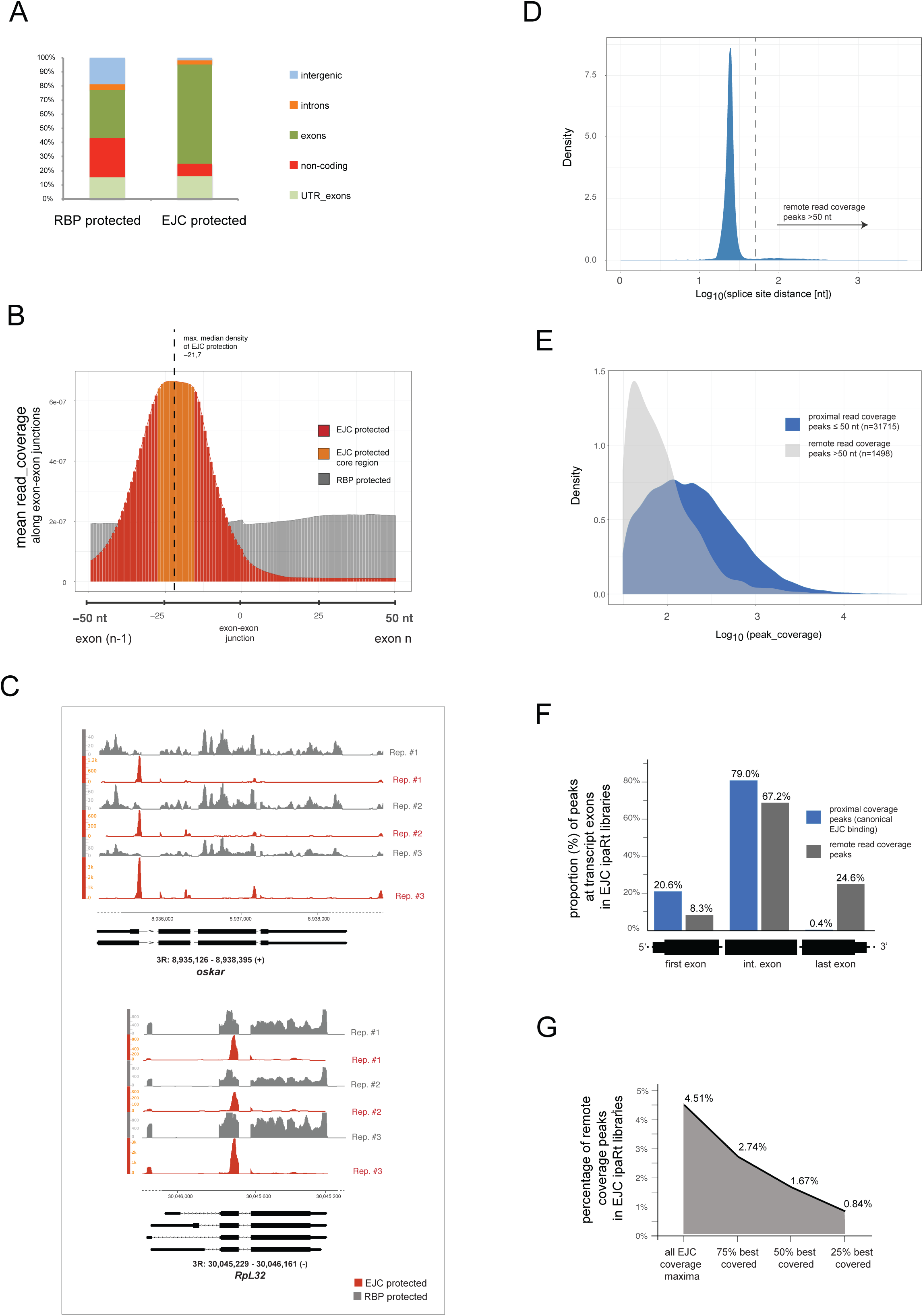
EJC binding to mRNA in *Drosophila* cytoplasm occurs within exons at canonical deposition sites. **(A) EJC ipaRt library reads map to exonic sites in the Drosophila genome.** Summary of genomic features detected in mRBP footprinting and EJC ipaRt sequencing results. Y-axis indicates proportion (percent) of all uniquely aligning read-counts. Color-code of genomic features is highlighted in the legend on right side of the plot. Note that EJC protected sites map in majority to exons and UTR-exons. **(B) EJC protection median is on upstream exons approximately −22 nt from the 3’ end.** Read coverage profiles from EJC ipaRt and mRBP footprinting cDNA libraries. Coordinates of metagene covering -+50 nt of exon-exon junctions are indicated on x-axis. Note position “0” defines the last nucleotide of upstream exons. Scale of mean sequencing read coverage for all normalized biological replicates is indicated on y-axis. Profiles from RBP protected and EJC protected junctions are presented in grey and red respectively. Note that mRBP footprinting sequencing reads show a homogenous RBP protection profile over the entire metagene body. On the contrary, sequencing reads from EJC ipaRt indicate a modal distribution of EJC protections with a median at 21.5 nt upstream of the exon-exon ligation point. Region of maximal EJC protection (protection core) spans from −27 nt to −15 nt and is highlighted in orange. **(C) Genomic coverage profile visualized by IGB viewer.** Coverage-profiles of 3 individual sequencing replicates from EJC ipaRt and mRNP footprinting along the *oskar and RpL32* genes. Size and position of gene regions are indicated. Exons and introns are indicated below the coverage profiles as boxes or lines, respectively. Coverage depth (normalized reads) is indicated on the y-axis. Note that cDNA reads from EJC protected fragments map upstream of and in direct proximity to splice sites, while reads from RBP protected fragments distribute over whole exon bodies. Sequencing coverage depth of EJC protected sites at exon-exon junctions is variable. **(D) Protection site peaks (modes) in EJC ipaRt libraries cluster primarily within 50nt around splice junctions.** Density plot highlighting distribution of coverage peaks in EJC ipaRt in respect to the splice site distance. Y-axis defines estimated density of peaks at defined splice site distances. X-axis indicates log10-transformed distance to splice site. Vertical dashed line highlights border between proximal and remote protection peak estimates. Note all peaks in ipaRt were defined within exons at a 20 nt frame resolution. Peaks chosen for the analysis were 2x more covered in ipaRt than in mRBP footprinting and had a coverage > 30 reads. **(E) Proximal (canonical) protection peaks in EJC ipaRt libraries are stronger covered than remote protection peaks.** Density plot showing the distribution of estimated coverage of splice junctions proximal (blue) and remote (grey) protection peaks. Y-axis defines estimated densities and x-axis the peak coverage. Note that peak coverage stands for the sum of all reads within a +/-10 nt window surrounding a protection site peak. **(F) Peaks within canonical EJC binding sites map to first and internal exons.** Bar plot showing proportion of peaks mapping to first, internal and last exons. Peaks mapping within or in direct proximity to canonical EJC binding regions (proximal peaks) are highlighted in dark blue. Peaks remote from EJC binding regions are highlighted in dark grey. Estimated peak proportions within exon classes are indicated in the plot. Y-axis indicates proportion in percent; X-axis indicates the tested exon classes of a metagene. Note that proximal peaks are nearly exclusively found in first and internal exons while remote peaks are detected in all three classes of exons. **(G) Proportion of remote protection peaks in EJC ipaRt libraries decreases with sequencing coverage.** Plot highlighting the proportion of remote peaks among all EJC ipaRt sequencing coverage peaks, and among sequencing coverage subsets of the best 75%, best 50% and best 25% covered protection peaks. Y-axis defines proportion (in %) of remote peaks. X-axis highlights peak coverage subsets. Note that proportion of EJC protection peaks reduces significantly when increased coverage cutoffs are chosen.

To define the median binding coordinates of the protected sites, we determined the sequence coverage +/−50 nt of exon-exon junctions in EJC ipaRt and mRNP footprinting (Figure 3B) and averaged the coverage profile over all junctions. The mean coverage profile in mRBP footprinting appeared evenly distributed, indicating an absence of protection bias (Figure 3B). In contrast, the protected sites in EJC ipaRt were located almost exclusively in upstream exons, with an EJC coverage median −21,7 nt 5’ to the exon-exon junction (Figure 3B), consistent with previous studies (4,16,44). Similarly, we estimated a 13nt long region of saturated RNA protection in the EJC RNA footprints, from −27 to −15nt 5’ of the exon-exon junction (Figure S3D) (4,8,36).

### Protection sites remote of exon-exon junctions in EJC ipaRt are not of EJC origin

Studies of mammalian EJCs have reported a high frequency of EJC-mediated protection outside of canonical binding sites (non-canonical EJC deposition sites) (36,37). To test whether this non-canonical distribution is representative of EJC protection across the *Drosophila* transcriptome, we determined coverage maxima for every exon-exon junction protected by EJC. The majority (∽ 95.5%) of protection peaks in EJC ipaRt libraries mapped to sites of canonical EJC binding, proximal to exon-exon junctions (Figure 3 C, D). The remaining ipaRt protection coverage peaks were located remotely (>50nt) of canonical EJC binding regions (Figure 3D,G and S4A).

Our analysis shows that proximal peaks map mainly to internal exons (79%), to first exons (20.6%), and only minimally to terminal exons (0.4%), as expected given the splicing-dependent deposition of EJCs upstream of splice junctions. Remote protection site peaks mostly mapped to internal exons (67%) and last exons (25%), and only minimally (8%) to first exons of the bound mRNAs (Figure 3F).

Three main features characterized the remote peaks in our EJC ipaRt libraries: low abundance, low sequencing coverage (Figure 3E,G) and relatively low expression compared with proximal peaks (Figure S4C,D). Further analysis using AME (45) revealed that the remote peaks in EJC ipaRt are significantly enriched in RNA binding motifs corresponding to known *Drosophila* splicing regulators (Figure S4E), whose binding might be a consequence of DSP crosslinking and co-purification due to direct interaction with the EJC. Our analysis shows that, in contrast to EJC in mammals (36,37), the proportion of remote EJC binding sites is negligible in *Drosophila*.

### EJCs mark multi-intron genes important for differentiation and development

We determined which RNAs are bound preferentially by EJC versus other RBPs using DESeq (46). This revealed a bias in EJC binding towards mRNAs of protein coding genes comprising greater than 1 exon (Figure S5D) and identified 3332 enriched and 4436 depleted genes (FDR < 0.05 and log2 fold change > log2(1.5)). Gene ontology (GO) term analysis of EJC enriched genes (Figure 4A) revealed significant association with genes involved in development or specialized cellular functions such as cell polarity,differentiation and cell signaling. In contrast, genes with homeostatic functions,involved in cytoskeletal and chromatin organization, in transcription or translation, and metabolic processes are biased for protection by other RBPs (Figure 4B, compare left and right plot). The bias of EJC binding to mRNAs from genes with functions in development and cell polarity, suggested that transcripts under spatial or temporal control might also show a preference for EJC binding. Consistent with this hypothesis, analysis of EJC and RBP protection sites on *Drosophila* transcripts annotated in the FlyFISH RNA localization database (47,48) revealed that localized maternal mRNAs are more likely to be bound by EJC than non-localizing transcripts (Figure 4C).

**Figure 4.**
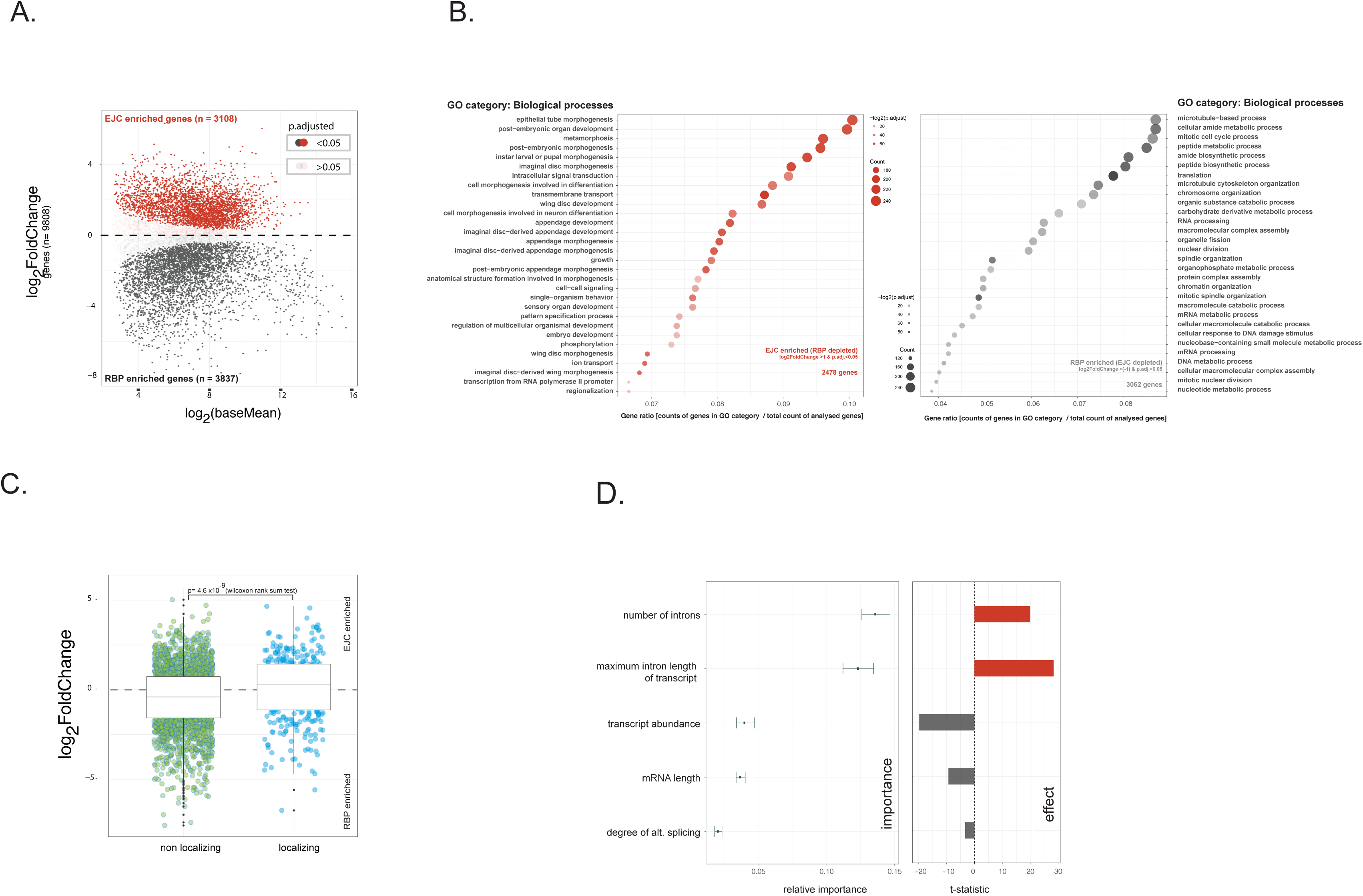
Preferential recruitment of EJC to mRNAs of genes with specialized cellular functions is determined by gene architecture. **(A) MA-plot of DESeq results from EJC ipaRt and mRBP footprinting.** Genes that are either enriched or depleted for EJC (RBP enriched) are indicated in red or grey, respectively. Genes that are not significantly different (p.adjusted >0.05) between the EJC ipaRt and mRBP libraries are transparent. Relative enrichment (log2FoldChange) is indicated on the y-axis. Base Mean of signal is highlighted on the x-axis. Dashed line defines log2foldChange =0. **(B) GO term analysis of EJC enriched and depleted transcripts |log2FoldChange| >1.** GO terms for biological processes of EJC enriched transcripts (left plot) and of EJC depleted transcripts (right plot) are presented to the left and the right of the plots, respectively. Y-axis list GO categories of the biological processes most highly represented. Gene counts in individual categories versus overall count of analyzed genes (Gene ratios) are shown on the x-axis. Legend indicating number of genes falling into a particular GO term (size of circles) and its significance of association (color of circles) are (circles) shown on the plots is highlighted in the center. **(C) Scatter-Boxplot showing DESeq enrichment estimates of *Drosophila* mRNAs with known localization patterns in the early embryo.** Gene products in DESeq analysis were subset to maternally expressed mRNAs, which localize or do not localize to specific foci in early embryos (48). Relative enrichment (log2FoldChange) is indicated on y-axis. Localization categories are indicated on x-axis. Non-localizing and localizing gene products are highlighted in green and blue, respectively. Horizontal solid line in corresponding box plots indicates median log2FoldChange for each of the categories. Dashed line highlights log2FoldChange =0. **(D) Result of Multiple regression model for EJC enrichment highlighting parameters which contribute most to preferential binding of EJC to mRNA.** Plots showing the relative contribution and effect of each factor (listed on y-axis) to EJC enrichment, as estimated from the full regression model. For the plot showing relative importance (left), estimates were obtained by bootstrapping the data; the dot indicates the median and whiskers indicate the 95% confidence interval. For the plot showing the effect of each factor (right), the T-statistic of each factor was calculated from the estimate and the standard error of each coefficient obtained after fitting the full model.

It was reported that in mammalian cells EJCs are enriched at mRNAs from alternatively spliced genes (38). In the fly, the EJC has been reported to promote correct splicing of long intron-containing genes (12,14). This relationship between EJCs and gene architecture led us to ask which features could explain the gene-to-gene variation in EJC deposition. We used a multiple regression model (see Materials and Methods) to assess how five features (number of introns, maximum intron length, transcript abundance, transcript length and the degree of alternative splicing) influence gene deposition of EJC. We checked that the effects estimated from the model holds given underlying correlations of the features. For example, after accounting for intron number, which has a strong effect on EJC binding (Figure S6A), we determined that intron length has a strong effect on EJC deposition (Figure S6B), while alternative splicing has only a minimal effect on EJC binding (Figure S6C). We assessed the relative importance of each feature from the multivariate regression analysis (Figure 4D). Two features dominated our model of EJC deposition: the number of introns per gene and the maximum intron length of the gene (Figure 4D) are positively associated with EJC deposition. The enhancement of EJC deposition with intron length agrees with previous studies showing that splicing of long intron-containing genes is sensitive to EJC activity (12,14).

### Splice site strength and hexamer composition influence EJC deposition

We estimated enrichment of EJC at the junction level using reads within +/-50 nt of the splice site and observed a strong dependency on the gene EJC estimate (Pearson correlation = 0.66, p <2e-16). This suggests that the junction’s EJC profile is primarily determined by its parent gene architecture. We next tested whether any exon-exon junction deviates significantly from its parent gene EJC binding. ∽31% of detected junctions have an enrichment that deviates significantly from the gene level (Figure 5A). Furthermore we observed that EJCs reads cover only a subset of junctions within a gene and show higher read-coverage coefficient of variation than reads protected by other RBPs (Figure S6D and S6E).

**Figure 5.**
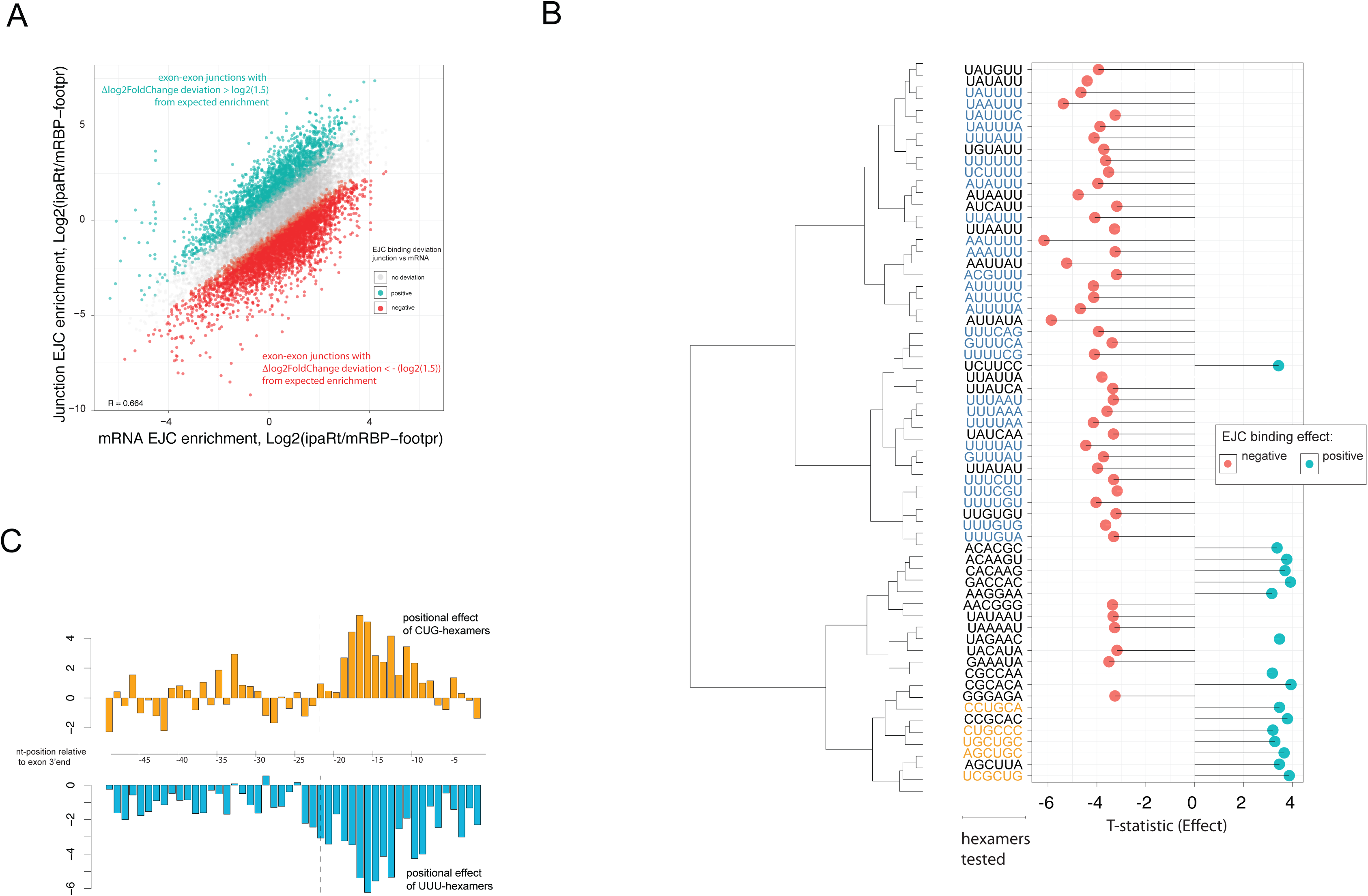
EJC assembly reveals a sequence bias. **(A) EJC enrichment – observed versus expected.** Scatterplot showing EJC enrichment defined by DESeq for individual exon-exon junction relative to DESeq estimates of corresponding templates. Enrichment for each exon-exon junction (log2FoldChange exon-exon junction) was calculated from read counts falling within 50bp upstream and 50bp downstream of the exon-exon junction. The EJC enrichment for the exon-exon junction was compared to its associated gene and a Z score and p-value was calculated. Exon-exon junctions whose EJC enrichment was not significantly different from that of its associated gene (p.adjusted >0.05) are shown in grey. Exon-exon junctions with significantly higher EJC enrichment than their associated genes are highlighted in red, and those with significantly lower EJC enrichment are highlighted in blue. The exon-exon junction EJC enrichment is highly correlated with the corresponding gene EJC enrichment (Spearman correlation = 0.664) and twice as many exon-exon junctions show significantly lower EJC enrichment than their corresponding gene product: 4334 vs. 2275, out of a total of 15186 junctions tested). **(B) Hexamers containing CUG and UUU contribute most to deviation between observed and expected EJC enrichment.** Plot showing the effect of each hexamer on EJC enrichment for each exon-exon junction, after accounting for splice strength and presence of an ESS. For each hexamer, the deviation (of the exon-exon junction EJC enrichment from its gene EJC enrichment) was regressed against (1) the presence / absence of hexamer, (2) splice strength, and (3) ESS counts. The x-axis shows the associated T-statistic obtained for the hexamer effect after fitting such a model for each individual hexamer. Dendrogram of hexamers was performed using hierarchical clustering with Ward’s method, based on pairwise Damerau–Levenshtein distance modelling between all hexamers. CUG containing hexamers show a concordant positive effect and UUU containing hexamers show a concordant negative effect on EJC enrichment. **(C) CUG and UUU effect EJC binding is position biased.** Plot showing the positional effect of the tri-nucleotides CUG and UUU. Similar to the analysis performed for hexamers, for each possible position of the tri-nucleotide, the deviation (of the exon-exon junction EJC enrichment from its gene EJC enrichment) was regressed against (1) the presence / absence of CUG / UUU at the position, (2) splice strength, and (3) ESS counts. Y-axis shows the associated T-statistic obtained for the position effect after fitting such a model for each individual position upstream of the exon-exon junction. X-axis defines the nt coordinates in the upstream exon. Both CUG and UUU show a highly positive or negative T-statistic around −18 bp upstream of exon-exon junction, indicating a strong position-specific effect.

We asked if a known sequence context related to splicing might be responsible for the variability in EJC binding to exon-exon junctions within a gene. We found that strong 5’ and 3’ splice site signals, and presence of intronic 5’ ISE correlates with increased EJC deposition, while the presence of an exonic 5’ ESS correlates with reduced EJC deposition (Table S1, Figure 7A), consistent with the fact that EJC assembly is dependent on splicing. Surprisingly, 5’ and 3’ ESEs and intronic ISEs at 3’ splice sites have no effect. Next we tested the effects of unannotated hexamers, while accounting for ESS, ISE and splice strength. We detected 63 hexamers that are associated with a change in EJC deposition. By clustering the hexamers based on sequence similarity, two major groups with either a negative effect or a positive effect on EJC assembly emerged (Figure 5B). Strikingly, 5 out of the 16 hexamers associated with increased EJC deposition contain the trinucleotide CUG, and 28 out of the 47 hexamers associated with decreased EJC deposition contain the trinucleotide UUU (Figure 5B). The strongest effect of these CUG or UUU trinucleotides is observed when they are present around the region downstream of EJC binding (approximately −16 to −18nt). This indicates that the sequence composition of this region is a strong determinant of EJC binding (Figure 5C).

### RNA structure modulates the degree and position of EJC binding in *Drosophila*

Deposition of an EJC at the first exon-exon junction and presence of a structured element next to the deposition site are required for localization of *oskar* mRNA at the posterior pole of the *Drosophila* oocyte (30,49). Interestingly, we observed a 2.38-fold stronger enrichment of the first *oskar* exon-exon junction than anticipated from *oskar* mRNA enrichment in our data, indicating that structures near EJC binding sites might affect EJC assembly.

To test whether RNA structures might have an impact on EJC binding, we estimated for every junction the probability of base pairing for each nucleotide −37 to +28 bp of the splice site. We observed three distinct average base pairing probability (bpp) profiles for exon-exon junctions with unaffected, positively correlated, or negatively correlated EJC binding (Figure 6A). Two regions showed significantly different bpps for junctions with a positive versus a negative effect on EJC binding (Figure 6B). The first region, located in the canonical EJC binding site, showed a decreased bpp for junctions with a positive EJC binding effect (Figure 6A,B) and increased bpp for junctions with negative EJC binding effect (Figure 6A,B). Surprisingly the second region, located directly downstream of the canonical deposition site (Figure 6A,B), showed an elevated bpp near junctions with positive EJC binding, but decreased bpp at junctions with negative EJC binding (Figure 6A). This result indicates that while EJC binding in *Drosophila* occurs on single stranded RNA (ssRNA), in agreement with previous reports (1,9), EJC binding to RNA may be enhanced by RNA secondary structures proximal to the EJC binding site.

**Figure 6.**
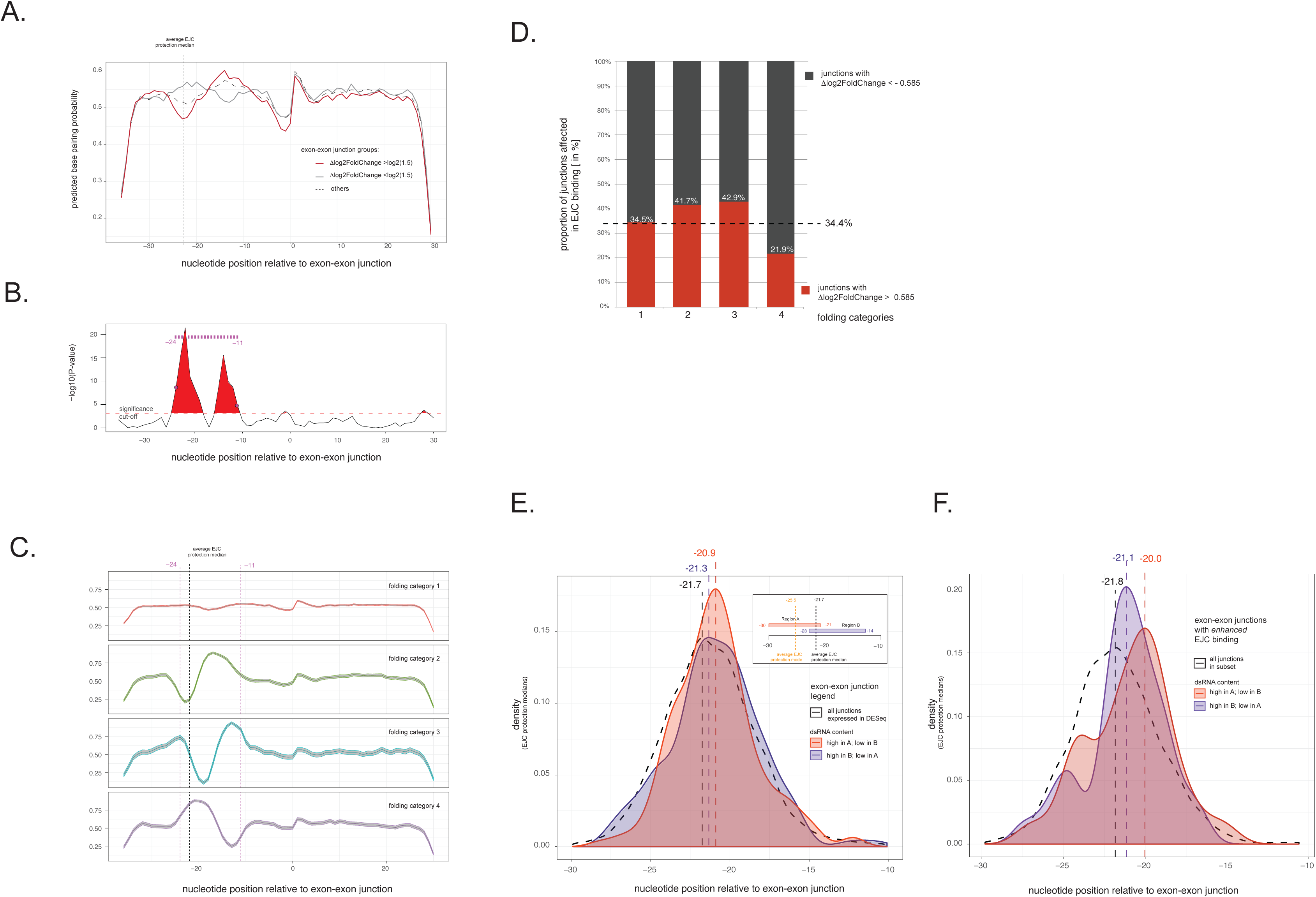
mRNA secondary structures modulate EJC binding. **(A) Pairing probability profiles of EJC bound exon-exon junctions.** Predicted Base pairing probability (bpp) profiles, by RNAplfold from Vienna RNA-package, of junctions with enhanced, inhibited and unaffected EJC binding. Black dashed, red and grey solid lines highlight average bpp profiles of junctions that are unaffected, enhanced and inhibited for EJC binding respectively. Note: The region utilized for RNA-Fold analysis covers the last 37nt of the upstream exon and the first 28nt of the downstream exon. Y-axis shows predicted bpp. X-axis highlights nucleotide positions relative to exonexon junction. Position 0 represents last RNA nucleotide of upstream exons. Dashed vertical line highlights the coordinate (−21.7) of the average EJC protection median. **(B) Definition of bpp regions distinct between exon-exon junctions with enhanced and inhibited EJC binding.** To define a cut-off for significantly distinct regions in the exon-exon junctions with enhanced and inhibited EJC binding, the bpp significance for each nucleotide in the two junction groups was tested by ANOVA, then subjected to permutation testing. Note that at alpha of 0.05 the regions −11 to −16 and −20 to −24 are significantly distinct in bpp between junctions enhanced and depleted in EJC binding. Y-axis defines – log10 transformed ANOVA p-values. Horizontal red dashed line indicates cutoff at a=0.05 of the permutation test. X-axis highlights nucleotide position relative to exon-exon junction. **(C) Folding categories of analyzed junctions**. To define folding categories, exon-exon junctions were clustered based on bpp estimates within the region 24 to −11 (see B). Clustering was performed with MClust using Gaussian mixture model assuming 4 clusters, and from the resulting categories the mean bpp profile of all cluster members are shown as individual bpp profile plots. Shaded region around solid lines indicates 95% confidence interval estimated using normal approximation to binomial distribution. Vertical dashed lines indicate borders of selected region for bpp profile clustering. Y-axis shows predicted bpp estimates, x-axis highlights nucleotide positions relative to exonexon junction. **(D) Impact of RNA structures on EJC binding**. Segmented bar plot showing proportion of significantly affected exon-exon junctions that are enhanced for EJC binding. Horizontal dashed line indicates overall proportion of junctions with enhanced EJC binding. Segments referring to proportion of junctions with enhanced or inhibited EJC binding are highlighted in red or grey, respectively. Y-axis defines proportion (in %) of all junctions significantly effected in EJC binding. X-axis indicates plotted folding categories (shown in panel C). Percent estimates for junctions with enhanced EJC binding are highlighted within the bar segments. Note that folding categories 2 and 3 have a positive, while folding category 4 - which has an increased bpp within EJC deposition sites -has a negative impact on EJC binding. **(E-F) RNA stem structures impact EJC binding coordinates.** Plots showing distribution densities of EJC protection site medians. Double stranded (ds) RNA content in Region A and B was identified using the mean of rounded bpp estimates [0 and 1] in the corresponding sequence stretches. High and low dsRNA content are defined as >0.7 and <0.3, respectively. The structural exonexon junction conditions tested, as well as the corresponding color-coding are indicated in the plot legend. RNA stem structural impact on EJC deposition coordinates in all junctions analyzed by DESeq is presented in **(E)**. RNA stem structural impact on EJC deposition coordinates in junctions with enhanced or inhibited EJC binding is highlighted in **(F)** Vertical dashed lines represent coordinates of estimated mean EJC protection site medians in each of the conditions. Y-axis defines EJC protection median density, x-axis indicates nt coordinates relative to the exon 3’end. Note that the strongest structural impact on EJC binding coordinates is present at exon-exon junctions with enhanced EJC binding.

Given the redundant information between bpp of each nt pair, we performed dimension reduction on bpp profiles within the −24 to −11 region (Materials and Methods, Figure 6B) using a Gaussian mixture model, to facilitate subsequent analysis of EJC binding. We obtained 4 folding categories (Figure 6C) and observed an association between significant positive EJC binding (log2fold-change >log2(1.5)) and junctions harboring folding categories 2 and 3, which contain a bpp elevation downstream of, or surrounding, EJC binding sites (Figure 6C, D). Junctions with an unstructured profile (folding category 1) show no such bias, and junctions with a bpp elevation in the EJC binding site (category 4) are associated with a negative EJC binding effect. This suggests that, when located near EJC deposition sites, RNA structures may positively affect EJC binding in *Drosophila* (Figure 6C, 6D lanes 2 and 3) and could explain the enhanced binding of EJC to the first exon-exon junction in *oskar* mRNA.

Previous in vitro experiments have shown that RNA secondary structures affect EJC assembly site coordinates (50). We asked whether predicted stable RNA stem structures within or flanking a canonical EJC binding site at exon-exon junctions have any impact on the precise coordinates of EJC assembly in the fly. For this we estimated potential dsRNA content in region A, spanning from −30 to −21 and for region B spanning from −23 to −14 within exon-exon junctions (see sketch in Figure 6E). We focused on junctions with a positive EJC binding effect to allow robust estimates based on junctions with high coverage. The analysis revealed a strong downstream shift of EJC assembly coordinates when dsRNA content in region A was high and low in region B, and only a minor shift when the dsRNA content was low in region A and high in region B (Figure 6F). Taken together, these observations confirm that the structural context of exon-exon junctions not only affects EJC binding efficiency, but also directs the site of EJC assembly.

### Comparison of human and *Drosophila* datasets reveals common factors that influence EJC enrichment

Previous studies of the EJC in *Drosophila* and human have reported differences between these two species in terms of EJC protein components and cellular function. We asked whether any of the factors we identified in *Drosophila* would also influence EJC deposition in human cell lines. Using the tools and models described in previous sections, we analyzed CLIP enrichment of the EJC component BTZ in HeLa cells (51). Results from the multiple regression analysis showed that the number of introns is a major determinant of the gene-to-gene variation in EJC deposition (Figure S8A). This agrees with our finding in Drosophila and suggests that this mechanism is conserved between *Drosophila* and humans. In contrast to our observations in the fly, we found that in mammals the extent of alternative splicing in genes has a small but positive effect on EJC deposition (Figure S8A), in agreement with previous findings that alternatively spliced genes are over-represented in EJC enriched genes (37,38). However, in contrast to *Drosophila,* in humans intron length did not facilitate but rather antagonizes EJC mRNA binding (Figure S8A).

Next we investigated the factors that determine variability in EJC deposition within genes (Table S2, Figure S8D). Similar to our *Drosophila* findings, high 5’ and 3’ splice strength and presence of an ISE at the 5’ junction enhances EJC deposition within a junction. In this analysis, presence of ESEs in the upstream, and of ESS in the downstream 50nt strongly affects EJC deposition, something we did not observe in *Drosophila* (Figure 7A). A possible explanation for this is that ESE/ESS/ISE sequences are annotated based on experiments in human cell lines and may not exert a similar effect in *Drosophila*. Among the 238 ESEs, we observed 28 hexamers containing AGAA which is similar to a motif (GAAGA) found in a previous EJC clip study (37,38). We separated ESEs according to presence of AGAA, and indeed found that the ESEs containing AGAA are associated with stronger EJC deposition. We asked whether bpp-profiles differ between junctions with positive and negative EJC binding. We found that an overall negative bpp (−24 to −18) around the EJC deposition site favors EJC deposition (Figure S8B,C). Unlike *Drosophila*, we did not find other regions whose bpp are associated with increased EJC deposition in mammalian cells. Taken together, these results indicate that across *Drosophila* and human, not only do conserved regulatory mechanisms such as intron counts, splice strength, or structural hindrance within EJC deposition sites influence EJC deposition, but also divergent regulatory factors such as ESEs in mammals or RNA folding in proximity of EJC deposition sites in *Drosophila* can affect EJC deposition.

**Figure 7.**
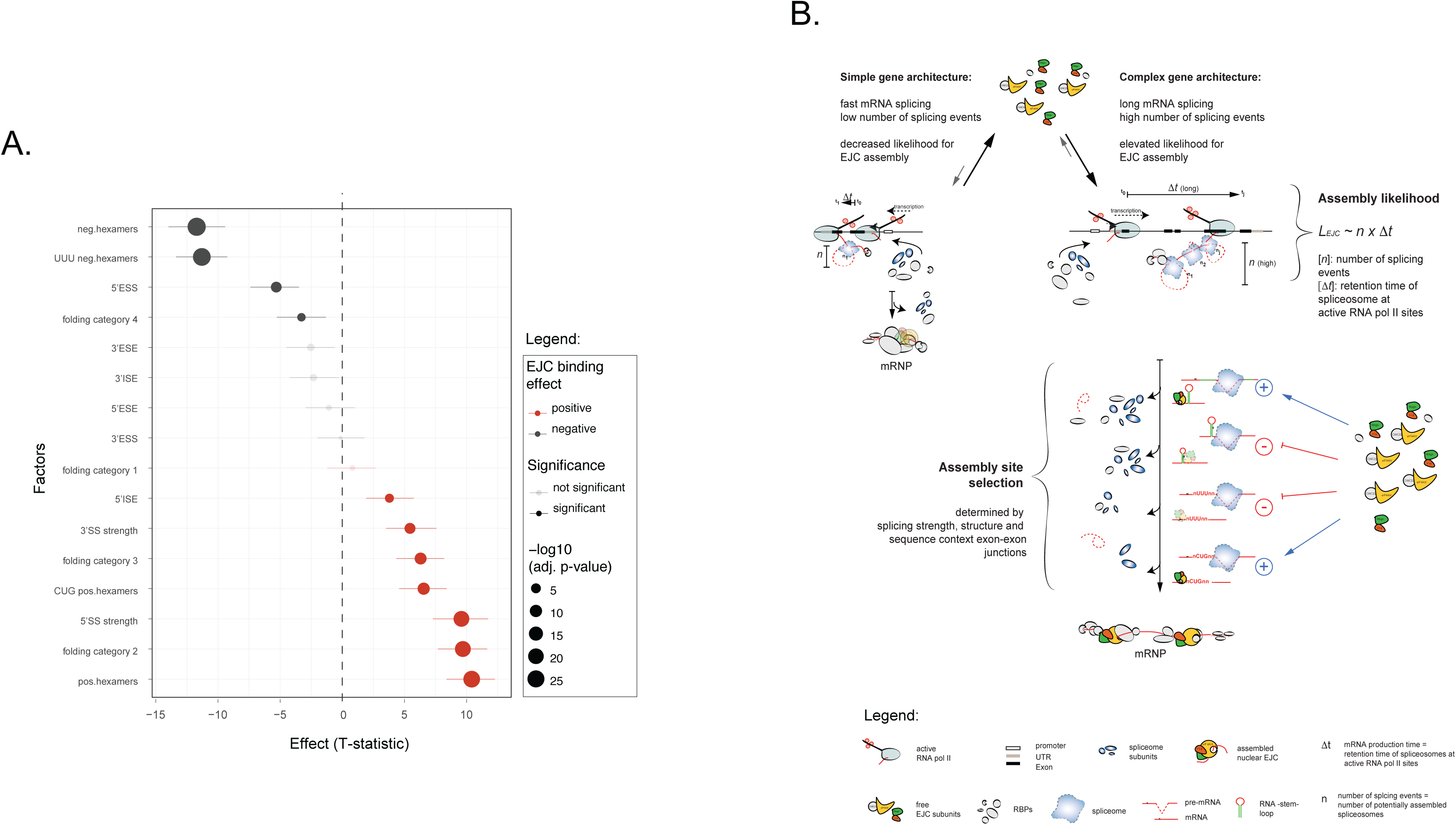
Summary of all identified factors regulating EJC assembly at exon-exon junctions in *Drosophila*. **(A) Factors in *Drosophila* that determine EJC binding variability within a transcript.** Plot comparing the effects of the difference variable on junction level EJC enrichment in *Drosophila*. For both datasets, we fitted a full linear model: Δlog2FoldChange ∽ splice + ESE + ESS + hexamer. Δlog2FoldChange is the difference between junction EJC enrichment and gene EJC enrichment. Splice, ESE and ESS are the splice site (SS) strength and number of ESE or ESS motifs present at 5”and 3”. The P value of each term was obtained using a t-test on each term (factor) and plotted. The effect of the term (sign of the coefficient) is reflected by the color of the bar. Red indicates a positive effect, meaning this variable has a positive effect on EJC deposition at the junction level, relative to the gene level. Y-axis indicates factors (terms) tested, x-axis highlights the result of the T-statistic. Summary legend is indicated to the right of the plot. **(B) Model of EJC assembly and variable binding along the gene’s transcript.** Co-transcriptional recruitment is affected by speed of mRNA production. mRNAs of simple genes with low complexity and low number of introns (n), are transcribed and processed faster than mRNAs from genes with diverged intron sizes and high number of introns. Longer processing increases retention time (Δt) of assembled spliceosomes at the site of transcription and thereby increases the likelihood of EJC assembly (LEJC ∽ n x Δt). Variable binding of EJC at mRNAs is a consequence of mRNA structure and sequence mediated hindrance or facilitation of EJC binding. RNA is highlighted in red, potentially pairing sites are highlighted in green. Note that binding of EJC is enhanced in proximity of dsRNA regions but repelled within dsRNA moieties. Legend is highlighted at the bottom of figure.

### Features that inform on EJC binding may predict mRNA localization

The bias of EJC binding to mRNAs from genes with functions in development and cell polarity suggested that EJC binding might be indicative of transcripts under spatial or temporal control. Consistent with this hypothesis, analysis of EJC and RBP protection sites on *Drosophila* transcripts annotated in the FlyFISH RNA localization database (47,48) revealed that localized maternal mRNAs are more bound by EJC than non-localizing transcripts (Figure 4C). We postulated that modalities underlying EJC enrichment at the gene and junction level can inform us about the localization of a transcript and applied decision tree learning on RNA localization using the R package rpart (52). Given the imbalance between localized to non-localized dataset, we used Cohen’s kappa coefficient to assess the predictive value of the model using different data groups. The use of gene features and features of the most enriched EJC junction is sufficient to achieve predictive accuracy comparable to a model incorporating all possible features (Figure 8A). Using the gene and highest EJC covered junction information as a working model, we asked which variables are important in this model. To prevent bias in estimate, the data were bootstrapped 1000 times. We observed that transcript length, maximum intron size in the gene, and 5’ splice strength and folding of the most enriched junction are more useful predictors (Figure 8B) compared to other variables. This suggests that features of gene architecture that orchestrate EJC deposition can distinguish localizing and non-localizing mRNA and might be important for mRNA localization.

**Figure 8.**
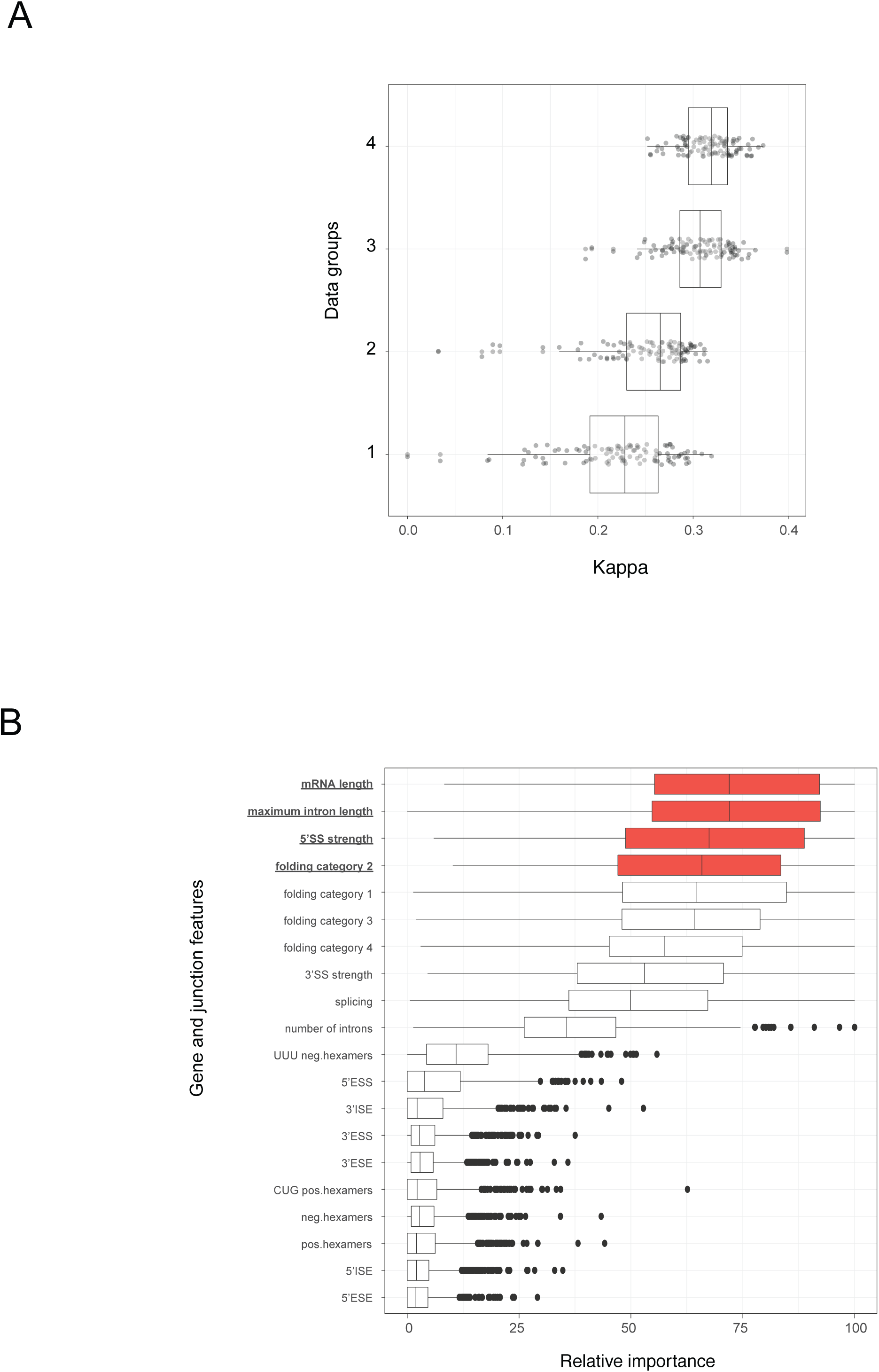
Gene and junction features inform on mRNA localization. **(A) Cohen’s kappa coefficient for different models.** Boxplots showing the distribution of Cohen’s kappa coefficient obtained after fitting tree-model for classification by recursive partitioning (over 100 bootstraps of 4 different data groups) using R package “rpart” (URL: http://CRAN.Rproject.org/package=rpart) (52). Group 1 includes gene features of the mRNA that influence EJC deposition from Figure 4E. Group 2 includes junction features (from Figure 7A) of the junction with the highest EJC deposition. Group 3 includes all variables in Group 1 and Group 2. Group 4 includes all the variables from Group 3 and EJC estimates (log2foldchange for the gene and highest EJC deposited junction, and the delta value). Median values are 0.246 (Data Group 1), 0.267 (Group 2), 0.306 (Group 3), 0.315 (Group 4). **(B) Relative variable importance for all features.** Boxplots showing the distribution of variable importance for all variables obtained after fitting tree model for classification, rpart over 1000 bootstraps of data group 3. The variable importance is an estimate of the usefulness of the variable in splitting the different classes in the regression tree. Highlighted in red are variables that are likely more useful in the classification, compared to the other variables.

## Discussion

### ipaRt: a method for high-confidence identification of protein binding sites on RNAs *in vivo*

We have profiled the landscape of EJC binding across the transcriptome of a whole animal, *Drosophila* melanogaster, and determined the parameters that influence the distribution of the complex on RNAs in the organism. Previous knowledge of EJC-RNA interactions was based on UV-crosslinking-based experiments in specific cell types grown as homogeneous cultures for the individual studies (37,38,43,53-60). While UV-crosslinking remains a method of choice for identification of protein binding sites on nucleic acids, due to the inefficient penetration of UV light into tissues and organisms, the method is most useful when applied to cells in culture. In contrast, our analysis of EJC distribution in the tissues of whole *Drosophila* flies was made possible by ipaRt, which employs the crosslinking agent DSP to freeze protein-protein interactions within otherwise dynamic RNP complexes, such as the EJC.

We have demonstrated that DSP-mediated covalent bond formation between the RNA helicase eIF4AIII and the Mago-Y14 heterodimer preserves EJCs in their “locked” state on mRNAs (1,2,8,9) and that efficient recovery of the bound RNAs does not require their crosslinking to eIF4AIII using UV light.Our “ipaRt” approach, like CLIP and iCLIP (53,56), enables highly stringent washing of the samples. In support of the robustness and reliability of our DSP based assays, we demonstrated high reproducibility not only among technical but also biological replicates of EJC ipaRt, as well as mRBP footprinting sequencing results (Figure S5A).

Furthermore, ipaRt allows enables use of non-RNA-binding subunits of the EJC, such as Mago, as immunoprecipitation baits. This is highly relevant in the context of studies of the EJC, as we and others have shown that its RNA-binding subunit, the RNA helicase eIF4AIII, may have other, EJC-independent functions in the cell (Figure S1C) (61,62). ipaRt afforded us the option of using Mago as our EJC bait, and indeed this is a main reason for the high-quality definition of the EJC binding landscape in the fly cytoplasm that we have achieved. The protection site reads we obtained from EJC ipaRt map almost exclusively to canonical EJC deposition sites (4) with a median protection ~22nt of the upstream exon’s 3’end. In contrast to mammalian EJC CLIP and RIP studies, in which eIF4AIII served as an immunoprecipitation bait (36,37), EJC ipaRt sequencing reads mapping to regions distant from canonical deposition sites are of low abundance and sequencing coverage. While this discrepancy could reflect differences in EJC engagement in humans and *Drosophila,* it more likely reflects the choice of bait or the cell compartment in which the analysis was executed. Indeed, a recent study in human cell lines revealed that when the cytoplasmic EJC component Btz was chosen as the immunoprecipitation bait rather than eIF4AIII, the proportion of non-canonical EJC deposition sites was negligible (38).

Finally, in ipaRt the DSP crosslinker is applied *ex vivo* during tissue disruption, and does not require inhibition of translation *in vivo*. We therefore consider ipaRt a method of choice for functional investigations of protein-RNA complexes in fully developed organisms and organ tissues.

### Regulation of EJC assembly in *Drosophila*

Through our analysis, we defined factors that contribute to or inhibit EJC assembly on mRNAs and at individual exon-exon junctions in *Drosophila*. From this we deduce that the landscape of EJC binding to RNAs is sculpted through regulation of EJC assembly at two levels in the fly (Figure 7B).

At the upstream regulatory level, the degree to which EJCs are assembled on an mRNA is dictated by the complexity of the gene’s architecture: mRNAs produced from genes of simple architecture are marked by fewer EJCs, while mRNAs from genes of complex architecture, comprising multiple splice sites and long introns are EJC bound to a higher degree. Due to the EJC’s assembly co-transcriptionally during splicing (4-7) it is not surprising that mRNAs from genes containing a larger number of introns are more likely to be bound by EJC. However, it is unprecedented that EJC assembly is enhanced on mRNAs of genes with large introns (Figure 4D). Although the EJC has been implicated to play a role in exon definition in the case of long-intron containing genes in *Drosophila* (12,14), the fact we observe a preference of EJC binding to the mRNAs, but not to the pre-mRNA splice junctions of long-intron containing genes (Figure S6F) disqualifies this simple explanation. We therefore propose that it is the number and the resting time of cotranscriptionally assembled spliceosomes at active RNA pol II sites that determines the likelihood of CWC22-dependent eIF4AIII recruitment and deposition, and thereby ascertains the degree of EJC assembly at mRNAs (Figure 7B).

At the downstream regulatory level, after EJC assembly rates at transcripts are defined, deposition of EJCs along mRNA exon-exon junctions is modulated by the structural and sequence context of the splice sites (Figure 7 A,B). dsRNA stem structures in exon-exon junctions of *Drosophila* mRNAs either antagonize EJC assembly when present within canonical EJC deposition sites, or enhance EJC assembly when located in the vicinity of the EJC deposition site (Figure 6D, 7A). Absence of dsRNA within the EJC binding moiety is in agreement with reported preference of EJCs for ssRNA (1,9,50); yet it remains to be elucidated, how and why EJC binding is positively affected when RNA-stem structures are found in its direct proximity on the bound template.

While it is likely that the structural context of exon-exon junctions in *Drosophila* directly influences the degree of EJC assembly, sequence composition derived effects on EJC binding to mRNA are the consequence of the assigned roles of these sequences during pre-mRNA splicing. We have demonstrated that exon-exon junctions with strong 5’ and strong 3’ splice sites (ss) are biased towards junctions with enhanced EJC binding (Table S1, Figure 7A). Importantly regulation of weak 5’ and 3’ ss which commonly occur at alternatively spliced junctions, *cis*-acting splicing regulatory elements (SREs) were shown to be of importance (63-65). In light of the negative impact of alternative splicing at the level of EJC mRNA binding (Figure 4D), it is not surprising that conventional ESEs and ESSs hardly affect EJC binding at the level of individual exon-exon junctions. However, whether the position-dependent bias mediated by the UUU-triplet and CUG-triplet containing hexamers towards inhibited or enhanced EJC binding which we have discovered in our *Drosophila* data set (Figure 5B, C) is due to a direct or indirect influence of these hexamers on splicing remains to be addressed. UUU-triplet containing hexamers which are strongly biased towards inhibition of EJC binding, could potentially function as yet undefined 5’ESS in *Drosophila*. Interestingly, CUG-triplet containing hexamers which are strongly biased towards enhanced EJC binding, share some sequence similarity with a previously predicted CUG containing 5’ ESE of short intron splice sites (64). It therefore appears likely that the CUG-triplet and UUU-triplet hexamers exert their effect on EJC binding as a yet undefined and novel class of SREs.

### RNA modalities influencing EJC binding in mammals and fly

In agreement with previous reports in mammals (36-38,66), we find that the extent of EJC occupancy varies between mRNAs and exon-exon junctions also in *Drosophila*. The splice site score next to a junction correlates with increased EJC deposition in the fly, and this relationship between splicing efficiency and EJC deposition has been proposed also in previous mammalian studies (67,68). Analysis of published mammalian Btz iCLIP data (38) revealed several modalities that correlate with the increased binding landscape of the EJC on mRNAs in mammals as well as in *Drosophila*, including the high number of introns, high transcript abundance and the sequence context of individual exon-exon junctions (Figure S8). Interestingly, the presence of long introns has a slightly negative effect, and the amount of alternative splicing a slightly positive effect on EJC occupancy in mammals (Figure 4D,S8); the latter is in agreement with previous observations (36-38). Previous studies on mammalian tissue cell cultures have reported that EJC enriched junctions contain a relatively high proportion of so called non-canonical protection sites, which were enriched for RBP consensus sequences of the SR protein family (36,37). Our analysis of Btz iCLIP data from mammals (38) confirms that the presence of ESEs in upstream exons and 5’ISEs in introns correlates with enhanced EJC binding (Table S2). Moreover, we have identified a group of junctions in mammals containing AGAA hexamers that are biased for enhanced EJC binding (Figure S7A), but their effects are not especially strong near the canonical EJC deposition site (Figure S7B). These hexamers match the AGAA encompassing consensus sequence of the mammalian SR protein SRSF10 known to function as splicing enhancers (69,70) and have been found previously in EJC bound exon-exon junctions (36-38). Not only do our *in silico* results agree with these reports and support the proposed cooperative binding of EJC with SR proteins (36-38), they also partially explain the EJC’s preference in mammals for alternatively spliced mRNAs.

One observation deriving from the analysis of published mammalian Btz iCLIP data sets is surprising, however. While we have observed junctions in *Drosophila* to be enhanced or inhibited in EJC binding by specific base pairing probability (bpp)-profiles, thus by specific RNA folding categories (Figure 6AD), we could not detect any striking differences between overall bpp-profiles of exon-exon junctions with enhanced or inhibited EJC binding in mammals (Figure S8B). Indeed, the only aspect of RNA structure shared by mammals and *Drosophila* is the negative effect of dsRNA when directly overlapping with the canonical EJC deposition site (Figure S8C) (50). *In Drosophila*, however,the presence of dsRNA close to canonical deposition sites enhances EJC binding, an effect that is not observed in mammalian cells.

### Insights into evolution and divergence of EJC functions

Our findings regarding the differences in the RNA modalities enriched at highly occupied mammalian and *Drosophila* EJC sites provide insight into the expansion of EJC functions during eukaryotic evolution. Spliceosome catalyzed splicing reactions are bidirectional and efficient formation of exon-exon junctions during RNA maturation is achieved by Prp22-induced release of spliceosomes from mRNAs (71-73). Importantly, EJC is absent in organisms with low rates of RNA splicing such as *Saccharomyces cerevisiae,* but present in organisms with high splicing rates such as *Schizosaccharomyces pombe* (74-76). This suggests that with the increased demand for splicing accuracy in higher eukaryotes, the EJC evolved to function as an exon-exon junction “lock” inhibiting spliceosome reassembly. Ensuring splicing reaction directionality, the EJC further evolved to become a central component of the NMD pathway (7,20-26) in mammals, where more than 95% of all genes are alternatively spliced (77). This may provide an explanation as to why EJCs in mammals are enriched on alternatively spliced transcripts. In *Drosophila*, where only 30% of all genes appear to be alternatively spliced (78), the EJC is not a component of the main NMD pathway (31). We therefore propose that although the EJC-NMD pathway evolved before the segregation of the proto-and deuterostome clades, it gained importance to complement the faux 3’UTR-NMD pathway during the evolution of vertebrates (79), where RNA surveillance and spatio-temporal control of gene expression are essential.

Similarly, the recruitment of EJC and interacting proteins upon splicing of mRNA to facilitate the localization of the mRNA so far seems exclusive to *Drosophila*. Two *Drosophila* specific features that modulate EJC binding, namely the presence of a large intron and secondary structure near the junction, are also predictive of mRNA localization. Although the precise strength of association between these features and mRNA localization remains to be verified with larger and more quantitative datasets, previous studies with the SOLE in *oskar* RNA have shown that RNA structure and EJC binding are indeed crucial for *oskar* mRNA localization (30).

## Acknowledgments

We thank Marco Blanchette, Matthias Hentze, Isabel Palacios, and Jean-Yves Roignant for their gifts of antibodies and fly stocks. We thank Sandra Müller, Alessandra Reversi and Anna Cyrklaff for technical assistance. We thank Vladimir Benes and the EMBL Gene Core Facility for their advice and service. We thank Mandy Rettel, Frank Stein and the EMBL Proteomics Core Facility, for their service and assistance with mass spectrometry analysis. We are grateful to members of the Ephrussi lab for discussions during the course of the work.

## Author Contributions

Conceived and designed wet lab experiments: AO. Performed the wet lab experiments: AO. Contributed reagents/materials/analysis tools: AO, LG, NH.Conceived and designed the *in silico* analyses: LG, NH, AO. Analyzed and interpreted the results: AO, LG, NH, AE. Contributed to the writing of the manuscript: AO, LG, NH, JU, AE.

